# Discriminating bacterial phenotypes at the population and single-cell level: a comparison of flow cytometry and Raman spectroscopy fingerprinting

**DOI:** 10.1101/545681

**Authors:** Cristina García-Timermans, Peter Rubbens, Jasmine Heyse, Frederiek-Maarten Kerckhof, Ruben Props, Andre G. Skirtach, Willem Waegeman, Nico Boon

## Abstract

**Background:** Investigating phenotypic heterogeneity can help to better understand and manage microbial communities. However, characterizing phenotypic heterogeneity remains a challenge, as there is no standardized analysis framework. Several optical tools are available, such as flow cytometry and Raman spectroscopy, which describe properties of the individual cell.

**Results:** In this work, we compare Raman spectroscopy and flow cytometry to study phenotypic heterogeneity in bacterial populations. The growth stages of three replicate *E. coli* populations were characterized using both technologies. Our findings show that flow cytometry detects and quantifies shifts in phenotypic heterogeneity at the population level due to its high-throughput nature. Raman spectroscopy, on the other hand, offers a much higher resolution at the single-cell level (i.e. more biochemical information is recorded). Therefore, it is capable of identifying distinct phenotypic populations in an automated way when coupled with analyses tailored towards single-cell data. In addition, it provides information about biomolecules that are present, which can be linked to cell functionality.

**Conclusions:** We propose a workflow to distinguish between bacterial phenotypic populations using Raman spectroscopy and validated this approach with an external dataset. We recommend to apply flow cytometry to quantify phenotypic heterogeneity at the population level, and Raman spectroscopy to perform a more in-depth analysis of heterogeneity at the single-cell level.

## Background

Single-cell phenotypic differences arise even in genetically identical cultures (Govers et al. 2017). A definition of a phenotypic population is an observed cellular state within a given genetic and environmental background. It arises due to epigenetic variations, stochastic gene expression, cellular age or oscillations such as the cell cycle. This is one of the strategies that bacteria use to adapt to a changing environment, as well as to divide the labour within the community (Avery 2006; Ackermann 2015).

Phenotypic heterogeneity in laboratory cultures is well-documented. For example, it has been studied in bacterial subpopulations that could tolerate antibiotics (known as ‘persisters’) (Dhar and McKinney 2007), in the production of cytotoxin K in *Bacillus cereus* (Ceuppens, Boon, and Uyttendaele 2013) or in the differential expression of flagellin in *Salmonella typhimurium* (Stewart et al. 2011). The challenge remains to find tools to measure and quantify this heterogeneity (i.e. phenotypic populations), in order to be able to link heterogeneity with bacterial functionality. This would allow to manage - and potentially steer – microbial communities in order to optimize bioprocesses.

Several tools are available for single-cell phenotyping (Davis and Isberg 2016). Imaging techniques are popular, but they require tagged bacterial cells or a probe to visualize the bacteria or the molecule of interest (Taniguchi et al. 2010; Anetzberger, Schell, and Jung 2012), making them less suitable to study environmental communities. There are other techniques that do not require a probe, such as intrinsic fluorescence (Georgakoudi et al. 2007) or the detection of autofluorescent NAD(P)H (Bhattacharjee et al. 2017). However, the amount of information that can be gathered is limited compared to other techniques, such as transcriptomics, flow cytometry or spectroscopy techniques. Single-cell transcriptomics are also an option for bacterial phenotyping, but a few hundred cells are needed and only about 15 - 25% of the expressed mRNAs can be detected (Tang, Lao, and Surani 2011). This analysis requires for bacteria to be lysed, and it was found in *E. coli* that a single cells’ protein and mRNA copy numbers are uncorrelated for any given gene (Taniguchi et al. 2010).

A more high-throughput option for single-cell analysis is flow cytometry, which can measure thousands of bacterial cells per second. Individual cells pass through a laser, after which detectors collect information on the scattered laser light (forward scatter, FSC, and side scatter, SSC) and on fluorescent emissions of specific probes (Davey and Kell, 1996). To detect bacteria, general nucleic acid stains (such as SYBR Green I or DAPI) can be used (Koch et al. 2018). This technique allows to quantify cells, and also to identify different phenotypes in bacterial populations. For example, this technique allowed Sanchez-Romero and Casadesus (2014) to find a differential expression of a GFP-tagged gene related to antibiotic resistance in a *Salmonella enterica* population, and Cronin and Wilkinson (2008) to detect a heterogeneous response of *Bacillus cereus* endospores to different heat treatments. Furthermore, the information derived from the FCM measurements can be transformed into a fingerprint, and used to calculate inter- and intra-species variations in bacterial populations (De Roy et al. 2012; Props et al. 2016; Koch et al. 2014). Flow cytometry can also be used for bioprocess monitoring, as it allows to quantify the number of cells present in a reactor, their viability and activity, as well as their membrane potential over time (Díaz et al. 2010). When this technique is coupled to cell sorting (also known as FACS, or FCM Activated Cell Sorting), a follow-up analysis on the subpopulations can be made. For example, by doing a proteomic analysis to link these phenotypes to a certain functionality (Jahn et al., 2013), to further culture the cells, or by doing single-cell microscopy analysis (Nebe-von-Caron et al., 2000).

Raman spectroscopy is another single-cell technology that has been proposed to study phenotypic heterogeneity. It does not require labelling and is non-destructive. The laser excites individual cells, which leads to inelastic scattering, which in turn is collected in the form of Raman spectra. It is less throughput compared to flow cytometry: even when enhancing the signal with metals (known as Surfaced-Enhanced Raman Spectroscopy or SERS), each cell takes 1-3 seconds to measure (Liu et al. 2016). The resulting spectrum contains biochemical information of the molecules that are present in the cell – e.g. lipids, carbohydrates, nucleic acids and proteins- and can be used to classify bacteria according to phylogeny (Goodacre et al., 1998). This information can be quantitative if an internal standard for the molecule(s) of interest is made -for example, (Cowcher, Xu, and Goodacre 2013) quantified the dipicolinate (DPA) biomarker for *Bacillus* spores; or (Samek et al. 2016) quantified polyhydroxyalkanoates produced by *Cupriavidus necator* H16)-. Raman spectroscopy can also be linked to cell sorting -known as RACS or Raman Activated Cell sorting-to further study phenotypes (Zhang 2015a).

Raman spectroscopy can be used for the monitoring of bioprocesses, as it can measure over time compounds present in the supernatant such as glucose, protein production or others (Lee et al. 2004), as well as some Raman reactive compounds present in the bacteria, such as chlorophylls, carotenoids and other pigments (Jehlička, Edwards, and Oren 2014). Although this technique is used for bacterial identification (Huang et al., 2010; Almarashi et al. 2012; Strola et al. 2014; Pahlow et al. 2015), the study of phenotypic heterogeneity by Raman spectroscopy bacterial fingerprint remains relatively little explored.

Both flow cytometry and Raman spectroscopy give rise to data that need specific pre-processing and analysis (O’Neill et al. 2013; Saeys, Gassen, and Lambrecht 2016; García-Timermans et al. 2018; Ryabchykov, Guo, and Bocklitz 2018). While microbial flow cytometry is rather limited in its phenotypic resolution (i.e., only a few properties are measured per cell), Raman spectroscopy characterizes many more biochemical properties of bacterial cells. It therefore requires analysis of high-dimensional data, which can be challenging, but it allows to characterize phenotypic heterogeneity at a much higher resolution.

In this work, we analysed bacterial cells from nine phenotypic populations –with a different growth stage and/or from a different replicate-using flow cytometry and Raman spectroscopy. We demonstrate how these populations can be automatically retrieved using data-specific algorithms. We also demonstrate how metabolic inference can be performed based on Raman spectroscopy in a data-driven way. Finally, the advantages and disadvantages of these tools for microbial phenotyping are discussed. We will motivate that, in its current form, microbial flow cytometry can be used to quantify phenotypic heterogeneity and describe community-level dynamics, while Raman spectroscopy can be applied to describe single-cell heterogeneity and possibly identify separated phenotypic populations. We include a recommendation for microbiologists on how to employ Raman spectroscopy and flow cytometry in future phenotyping studies.

## Results

Throughout this paper, we define a ‘phenotypic population’ as a group of bacteria grown under the same environmental conditions (i.e. cells from the same biological replicate at a certain growth stage). This population will share morphological and/or metabolic traits that can be detected by flow cytometry and Raman spectroscopy. Samples of *E. coli DSM 2092* were measured in the lag, log and stationary phase. For every condition, triplicates of the cell culture were made. Thus, we expected to retrieve 9 phenotypic populations. As it will be argued in the discussion, this is not to say that there might not be other subpopulations in each of these ‘phenotypic populations’.

### Flow cytometry

Three biological replicates of *Escherichia coli* DSM 2092 were measured in the lag, log and stationary phase through flow cytometry (Fig. S1). Data was analysed at two levels: (a) the single-cell level (i.e., cells were analysed as individual instances) (b) the cell population level (i.e., cytometric fingerprints were constructed to describe population dynamics) (Fig. 1). t-distributed stochastic neighbourhood embedding (t-SNE) and principal component analysis (PCA) were used to visualize the data at the single-cell level (Fig. 1a-d, Fig. S3). Principal coordinate analysis (PCoA) was applied to visualize the differences of the phenotypic populations based on Bray-Curtis dissimilarities (Fig. 1f). As a validation, t-SNE was performed on the population level as well (Fig. 1e).

**Fig. 1:**
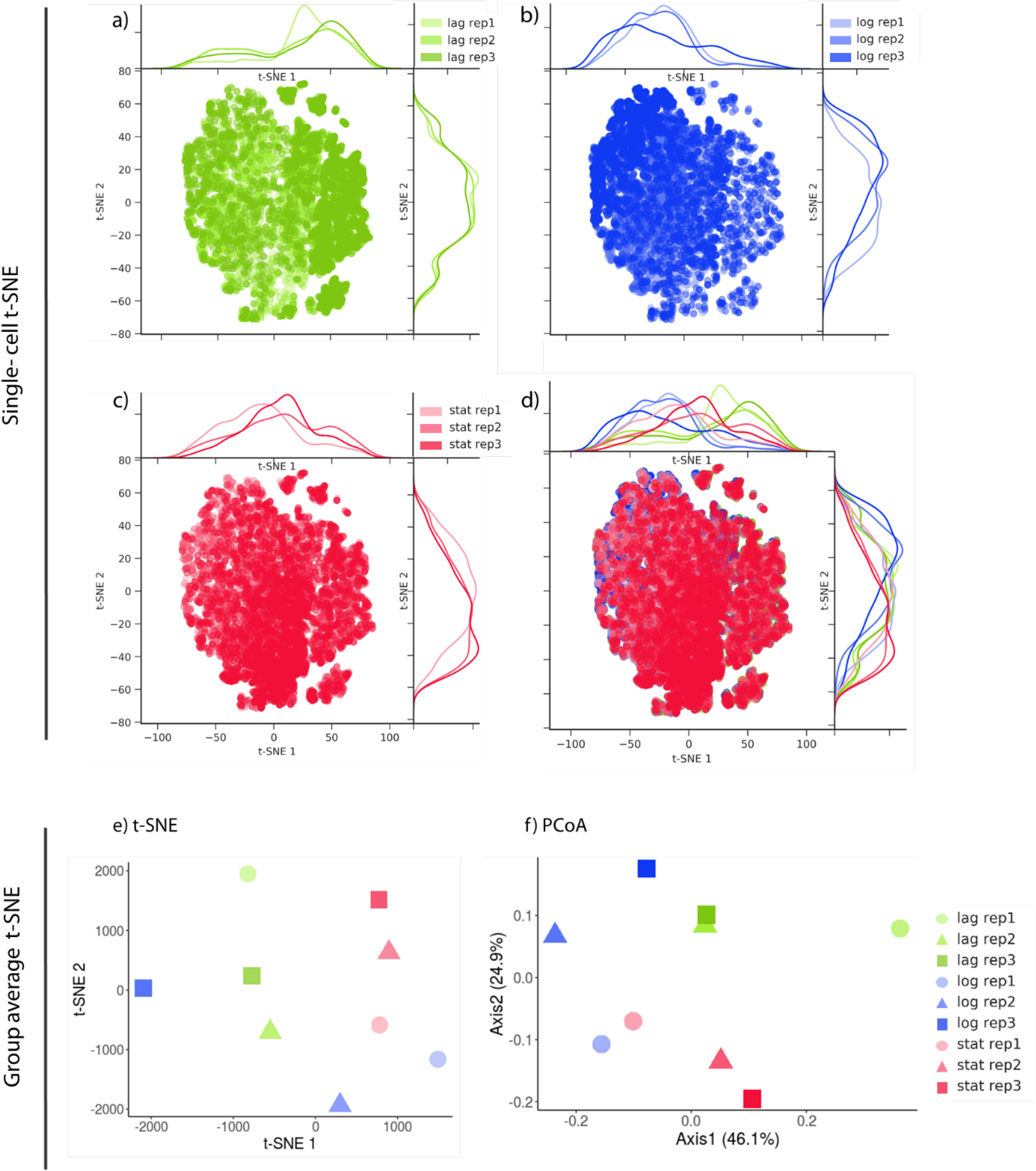
*E. coli* measured with flow cytometry and analyzed at the single cell level (a-d) and population level (e and f). t-SNE was performed on the aggregation of all samples (d), and visualized separatedly for each growth phase, to allow for easier interpretation (a-c). Distributions on the side represent the t-SNE distributions separately visualised for each growth phase/replicate to allow for easier interpretation. (e-f) Visualization of cytometric fingerprints at the sample level, using t-SNE (e) or PCoA (f).

No separated subpopulations could be distinghuished based on cytometric single-cell data (Fig. 1a-d). Yet, shifts in the distribution of cells were clear, both between different growth phases and replicates, as can be seen from the marginal distributions. Therefore, by creating cytometric fingerprints, which are vectorizations of the cell counts per bin, these differences could be quantified and visualized at the community level (Fig. 1e-f). Differences between fingerprints were calculated using the Bray-Curtis dissimilarity. Average dissimilarities per growth phase and replicate were summarized in Table 1. The average Bray-Curtis dissimilarity between samples within the same growth phase is smaller compared to samples that originated from the same replicate (Table 1). The lag phase for replicate 1 was quite different from the other samples (Fig. 1f).

**Table 1:** Average Bray-Curtis distance between the samples based on their growth phase or their replicate. A.U.= arbitrary units.

|  | **Average distance between samples (A.U., n=3)** | **Standard deviation (n=3)** |
| --- | --- | --- |
| Lag phase | 0.33 | 0.08 |
| Log phase | 0.33 | 0.06 |
| Stationary phase | 0.21 | 0.06 |
| Replicate 1 | 0.41 | 0.18 |
| Replicate 2 | 0.35 | 0.04 |
| Replicate 3 | 0.35 | 0.07 |

### Raman spectroscopy – clustering results

The samples used for flow cytometric analysis were fixed and analysed using label-free Raman spectroscopy following the protocol from (García-Timermans et al. 2018). To identify phenotypic populations, two clustering methods were used. First, using an agglomerative hierarchical clustering approach and secondly, using the PhenoGraph algorithm - a tool developed for the analysis of high-dimensional cytometry data. To determine the hierarchical clustering, the spectral contrast angle between samples was calculated (a measure of the spectra similarity). Then, phenotypic populations can be delineated by setting a threshold upon inspection of the resulting dendrogram after hierarchical clustering (Fig. 2). On the other hand, PhenoGraph makes use of *k*-nearest-neighbours clustering, in order to determine groups of similar cells, and as such, phenotypic populations. In other words, *k* expresses the amount of local information that is included when cells are grouped according to similar spectra. *k* will therefore, in a similar way as the threshold used in hierarchical clustering, impact the number of phenotypic populations that are defined (Fig. 3).

**Fig. 2:**
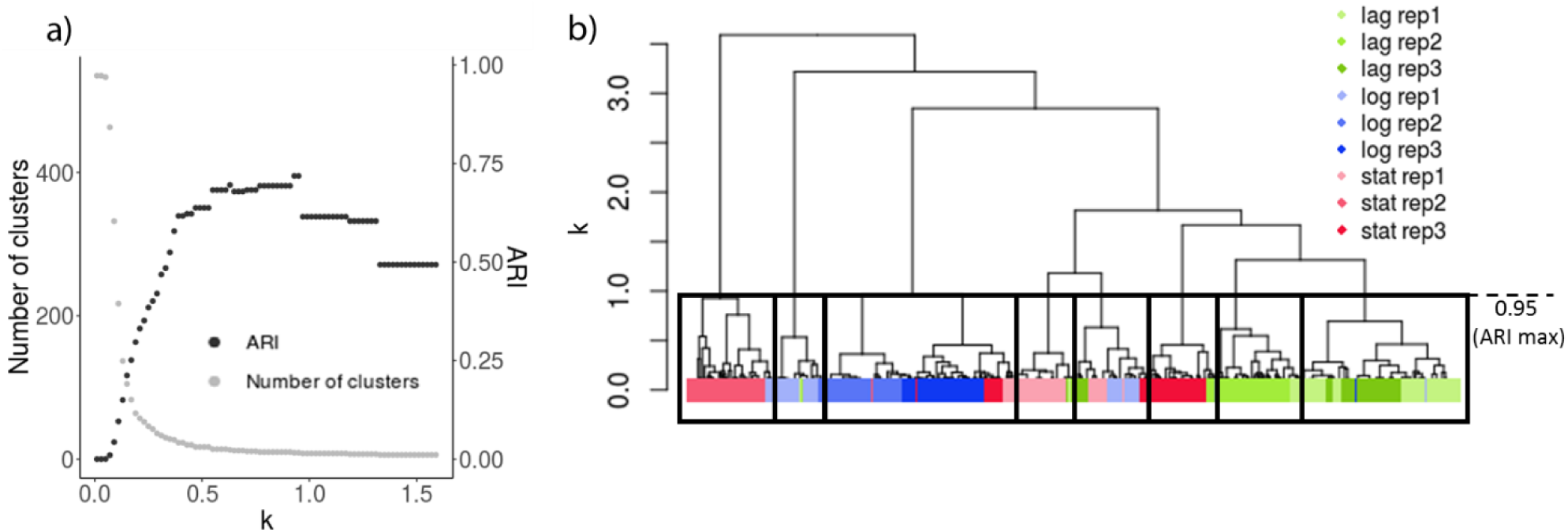
Hierarchical clustering of Raman spectra derived from the *E. coli* culture. Cells were measured in the log, lag and stationary phase using Raman spectroscopy. (a) Left axis, grey: Visualization of the number of clusters we can obtain by cutting the cluster at different heights (k). Right axis, black: ARI, which quantifies how many cells are identified as the expected phenotypic population when clusters are made at different levels. (b) Dendrogram representing the distance (calculated as the spectral contrast angle) amongst the Raman spectra. The black boxes dived the phenotypic populations when the dendrogram is cut at k=0.9. This is the maximum adjusted Rand index (ARI), as calculated in Fig. 2a.

**Fig. 3:**
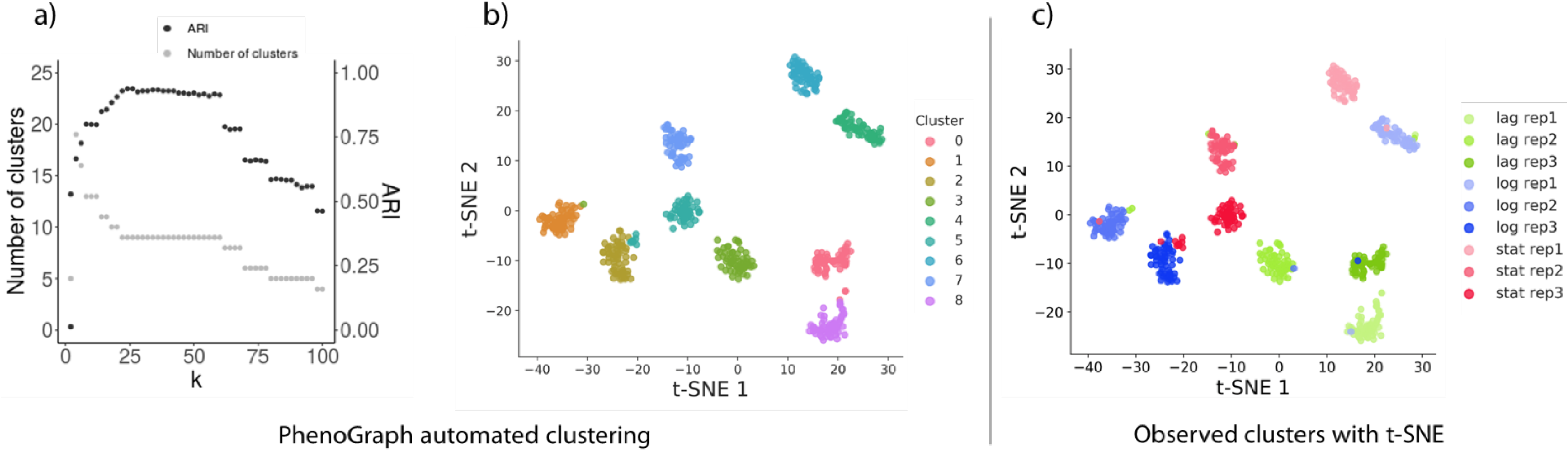
PhenoGraph clustering of Raman spectra derived from the *E. coli* culture. a) Influence of hyperparameter *k* on the automated identification of phenotypic populations. Left axis, grey: Visualization of the number of clusters in function of k. Right axis, black: Adjusted Rand index (ARI), which quantifies how many cells are identified as the expected phenotypic population when clusters are made at different levels (k). b) t-SNE visualization, colored according to PhenoGraph clustering with optimal ARI (*k*=22). c) t-SNE results colored according to growth phases and replicates.

The adjusted Rand index (ARI) was used to quantify similarity between the clusters that were determined by hierarchical clustering and Phenograph and the known phenotypic populations (i.e. growth phase and replicate). An ARI of 1 indicates perfect grouping of the data. The PhenoGraph algorithm resulted in a higher ARI as opposed to hierarchical clustering based on the spectral contrast angle (Fig. 2a vs. Fig. 3a). Inspecting the PhenoGraph results, there is a stable region for *k* that retrieves clustering according to both growth phase and replicate (i.e., nine clusters were found for *k* = 20,…, 60). A value of *k* = 24 or 26 gave rise to an optimal clustering (Fig. 3a). Smaller *k* allowed to inspect phenotypic populations at smaller scales and investigate the heterogeneity accordingly. See for example the clustering results for *k* = 15, which resulted in eleven different groups of cells (Fig. S5). Additional clusters that emerged were the result of splitting two clusters into two smaller ones. Likewise, larger *k* will result in larger clusters. For example, for *k* = 100, data is grouped in five clusters (Fig. S5). Structure in the data is retained, as clusters are merged either according to growth phase (clusters 0, 2 and 3) or replicate (cluster 1).

Often a single-cell was classified as the expected replicate, but in another growth phase (Fig. 3c). The samples in the lag phase seem to have a single-cell that is already in the log phase, and in the cultures in the log phase, we find a cell in the stationary phase (in replicate 3) and one cell in the lag and in the stationary phase (in replicates 1 and 2). It is also worth noting that some cells from replicate three seem to be between the log and the stationary phase.

### Raman spectroscopy – tentative region assignment

The Boruta algorithm, a variable selection algorithm based on Random Forests, was used to associate the most distinctive regions in the Raman spectrum with cluster assignments according to the hierarchical clustering and PhenoGraph algorithm. The cluster labels that resulted in an optimal ARI were used. Regions were linked with different molecules based on a recent summary from Wang et al., 2016. In this way, metabolic associations could be inferred that contained predictive power as a function of different phenotypic populations (Table 2).

**Table 2:** Tentative assignment of Raman spectra using the Boruta algorithm based on phenotypic identification using hierarchical clustering and PhenoGraph. The 10 highest ranked areas are shown. When there is no known compound in the spectral region, either the closest compound or a blank is shown. A.U.=arbitrary units.

| Wavelength<br>(cm<sup>-1</sup>) | Features importance |  | Tentative assignment |
| --- | --- | --- | --- |
|  | PhenoGraph (k=22,<br>9 clusters) (A.U.) | hclust (h=1, 8<br>clusters) (A.U.) |  |
| 1042 | 9.8 | 9.8 | Carbohydrates, Proline (1043) |
| 971 | 9.5 | 9.7 | v(C-C) wagging (971) // Phosphate monoester groups of phosphorylated proteins and cellular nucleic acids (970) |
| 945 | 9 | 8.9 | vs (CH<sub>3</sub>) of proteins (α-helix) (951) |
| 1057 | 8.9 | 8.8 | Lipids (1057) // Carbohydrates (1030-1130) |
| 1294 | 8.5 | 8.5 | CH<sub>2</sub> deformation (1295) |
| 1050 | 8.5 | 8.6 | Nucleic acids, CO stretching; protein, C-N stretching, PHB (1054) // Carbohydrates (1030-1130) |
| 1053 | 8.4 | 8.4 | Nucleic acid (1054) // Carbohydrates (1030-1130) |
| 1127 | 8.3 | 8.4 | v(C-N), protein (1127) // Carbohydrates (1030-1130) |
| 1046 | 8.1 | 8.2 | v<sub>3</sub>PO<sub>4</sub> 3- (symmetric stretching vibration of v<sub>3</sub>PO<sub>4</sub>3- of HA) (1044) |
| 641 | 8 | 7.9 | C-C twisting of tyrosine (642) |

To understand how the molecules in Table 1 vary from one group to another, the distribution of intensities of these Raman regions was plotted for every growth phase (Fig. 4). A Wilcoxon rank sum test with a Benjamini–Hochberg correction was performed for the ten highest ranked variabels according to the Boruta algorithm (Fig. 4).

**Fig. 4:**
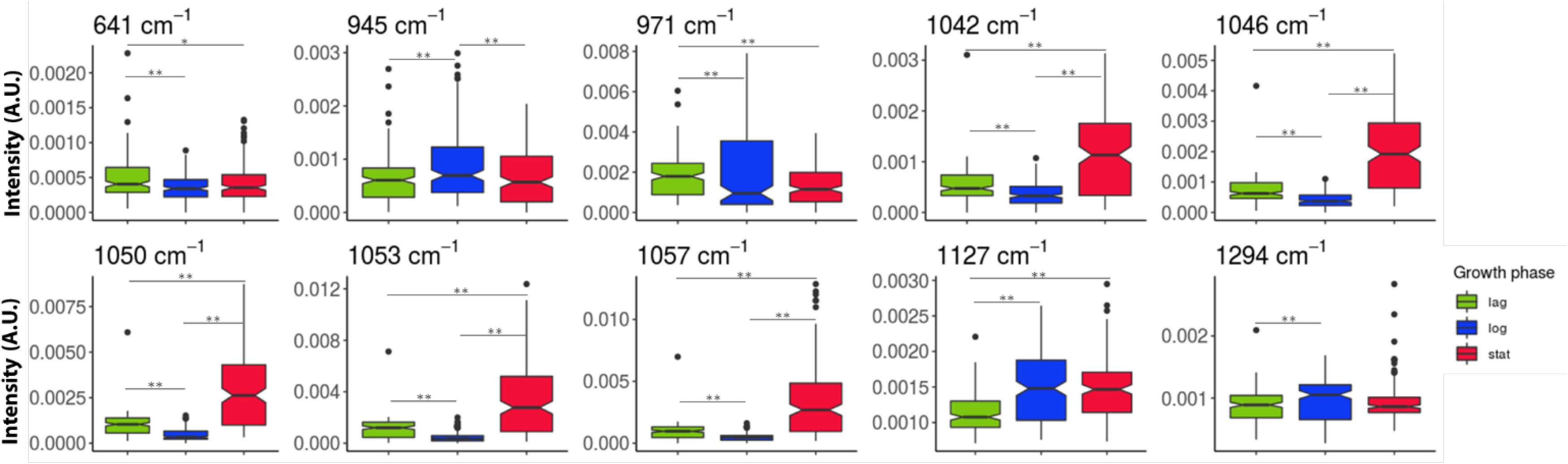
Distribution of intensities of the most relevant regions associated with phenotypic populations according to the Boruta algorithm. Boxplots represent the growth phases (replicates were pooled together). Groups were made according to the growth phase: lag (green), logarithmic (blue) or stationary (red). For every spectral region, a Wilcoxon rank sum test was made, with a Benjamini-Hochberg correction (upon rejection of the null hypothesis). Groups with significantly different peaks are signalled with (*) (p<0.05) or (**) (p<0.01).

To better understand what regions of the Raman spectra (and therefore, what biomolecules) were making these phenotypic populations different, we defined phenotypic populations at different levels (changing the *k* parameter in the PhenoGraph algorithm) and then used the Boruta algorithm to identify the most relevant regions.

When more phenotypic populations were distinguished (i.e. setting the value of k lower) more regions in the Raman spectrum were associated with differences in phenotypic populations. As shown in Figure 5, to find the phenotypic populations with different growth phases, 59% of the regions are included (green); to find the biological replicates, it is 67%; and to find both categories, 77%. Although this result was expected, a large number of Raman regions (48%) were relevant for all levels of classification.

**Fig. 5:**
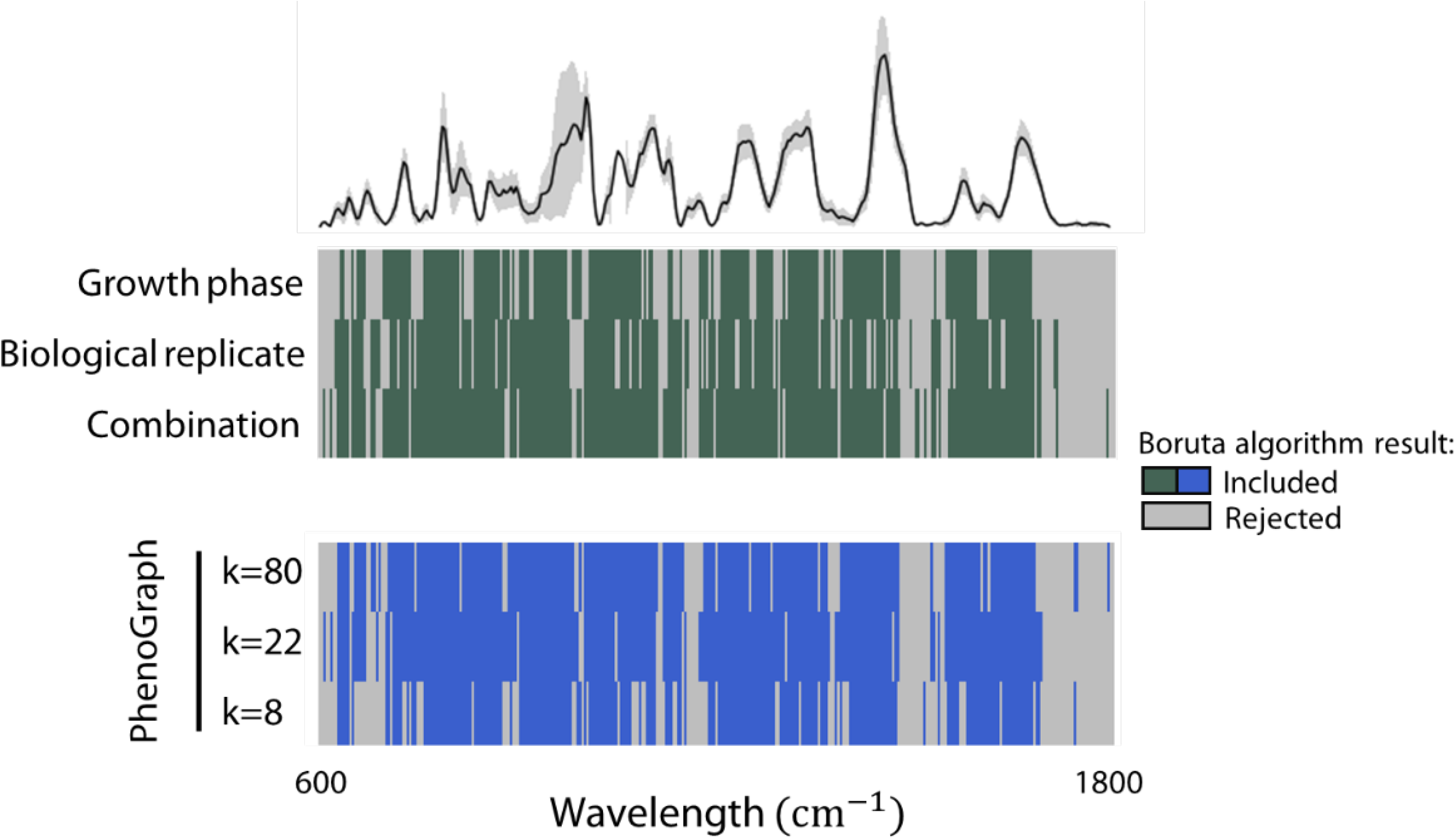
Spectral regions relevant for phenotypic classification at different levels, according to the Boruta algorithm. The top heatmap shows the results for the growth phase and replicates. At the bottom, the results for PhenoGraph with the clustering hyperparameter k as 80, 22 or 8 are shown. In green and blue, spectral regions confirmed by the Boruta algorithm as relevant, in grey the rejected. The average of all spectra is also plotted; the grey areas in the average spectrum correspond to the standard deviation.

### Validation of single-cell analysis of Raman spectra

To validate our workflow to analyse microbial single-cell Raman data, the dataset from Teng et al., 2016 was used. In this work, *E. coli* was exposed to different chemicals (ethanol, antibiotics, n-butanol or heavy metals) and the spectra of the bacteria were measured at several time points after the treatment (5, 10, 20, 30 and 60 min, 3h and 5h). Three replicates of the cell culture were made for each treatment. Here we show the results for cells treated with ethanol (Fig. 6), representative for what is observed in the other groups (Fig. S6). t-SNE was able to visualize groups of bacteria that received different treatments at different points in time. Furthermore, two subpopulations are seen in every group. They correspond to the replicates, where two replicate samples are separated, and the third replicate is either assigned to one of the two or divided amongst the two subpopulations. The optimal ARI is lower than the one reported for our own work, but still considerably higher than zero. This means that although the clusters assigned according to PhenoGraph have a better match with the treatments induced in our own dataset compared to this one, the clustering is still meaningful.

**Fig. 6:**
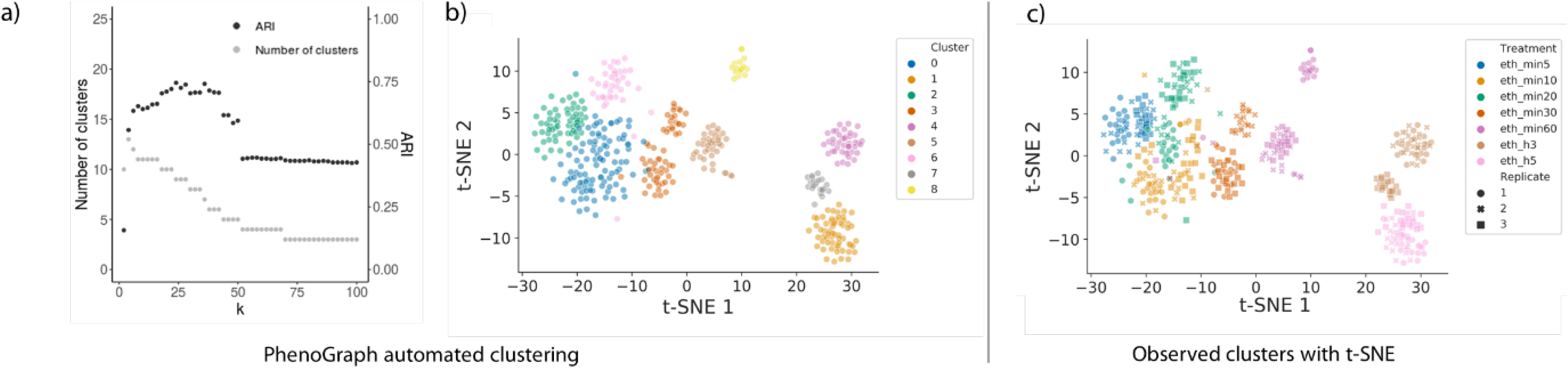
External dataset clustered using t-SNE. *E.coli* treated with ethanol and measured at time points 5, 10, 20, 30 and 60 min, 3h and 5h. (a) Influence of hyperparameter *k* on the automated identification of clusters. Left axis, grey: Visualization of the number of clusters in function of k. Right axis, black: Adjusted Rand index (ARI), which quantifies how many cells are identified as the expected phenotype when clusters are made at different levels (k). (b) Using the maximum ARI, samples were automatically clustered using PhenoGraph. (c) t-SNE was performed on the data set and samples were labelled according to their treatment. The shapes represent the sample replicate.

## Discussion

### Flow cytometry quantifies population shifts

When using single-cell data from flow cytometry, the different phenotypic populations overlapped and did not form separate clusters, as shown by both the t-distributed stochastic neighbourhood embedding (t-SNE) and the principal component analysis (PCA) (Fig. 1a, SI Fig. 2). However, in the t-SNE plot, a consistent shift in the cells distribution could be observed in response to the different growth phases (Fig. 1a-d). In other words, gradual shifts in the structure of the phenotypic population, i.e. the phenotypic heterogeneity, could still be detected, although individual cells could not be separated according to growth phase or replicate. As flow cytometry is capable of rapidly measuring a large amount of cells, the differences in phenotypic heterogeneity at the population level could be described by constructing cytometric fingerprints (Koch et al. 2014; Props et al. 2016). The Bray-Curtis dissimilarity was used to quantify these differences at a sample level. The average Bray-Curtis dissimilarity showed that the effect of the replicates exceeded the effect of the growth phase (except for the lag phase). This implies that the differences of *E. coli* cells in different growth stages are comparable or smaller to those the differences amongst replicates. The results from (Teng et al. 2016) analysed in Figure 6 leads us to hypothesize that when the community is steered with a stressor, the effect in the fingerprint is bigger, making the replicate effect smaller.

Flow cytometry is a high-throughput technique, able to rapidly measure hundreds to thousands of individual cells. By applying fingerprinting approaches to cytometry data, differences between microbial populations at the population level can be assessed and quantified. In this work, gradual shifts could be detected in the flow cytometric data at the level of individual cells, while at the sample-level (i.e. the population distribution level), differences between communities could be quantified (e.g., using the Bray-Curtis dissimilarity) and separated accordingly. In other words, while current resolution at the single-cell level appears to be limited for microbial flow cytometry, due to its high-throughput nature, its power lies in the possibility to characterize and phenotypic heterogeneity at the population level and quantify shifts in phenotypic heterogeneity. In this work, gradual shifts could be detected in the flow cytometric data at the level of individual cells, while at the sample-level (i.e. the population distribution level), differences between communities could be quantified (e.g., using the Bray-Curtis dissimilarity) and separated accordingly.

It is worth noting that in this work the effect of using additional or more specific labels for cytometric analysis has not been explored, which might improve the resolution. It is possible to add stains to target specific substrates (see the review of Léonard et al. (2016) on the use of individual and double stains, and an example of a three-color analysis by (Barbesti et al. 2000)), but the number of markers describing microbial cells using flow cytometry will never be of the same order as that of Raman spectroscopy. In eukaryotic flow cytometry, where the tagging of specific antibodies is much more feasible, 19-parameter flow cytometry is routinely used (17 fluorescence and two scatters) (Perfetto, Chattopadhyay, and Roederer 2004) and 30-parameter flow cytometry has just recently been published (Mair and Prlic 2018). However, the dimensionality of cytometry data in these settings is still much lower than the number of variables derived from Raman spectroscopy. Even in the best-case scenario, the dimensionality of flow cytometry data cannot get close to the number of parameters that Raman spectroscopy exhibits. On the other hand, depending on the research question, a high-dimensional tool might not be needed. For example, biomolecules that are associated with a phenotypic population might be known, and there could be a dye available to highlight these molecules. In this case, flow cytometry could be more relevant for phenotyping than Raman spectroscopy, provided the proper parameters are chosen to differentiate among treatments. In this work, we compared phenotypic populations that are not differentiated by (a) specific, known molecule(s) but used a general marker to characterize the DNA content. This explains why flow cytometry did not have enough resolution to differentiate the phenotypic populations.

### Raman spectroscopy detects phenotypic populations at the single-cell level

Raman spectroscopy is lower throughput for single-cell analysis when compared to flow cytometry, but it is able to retrieve much more information per cell. Its resolution is enough to conduct research at the single-cell level. The study of bacterial phenotypes using Raman spectroscopy has been done by other groups: to identify different growth stages in *L.casei* (Ren et al., 2017), stress-induced phenotypic populations (Teng et al. 2016), bacterial phenotypes with different antibiotic responses (REFAtha 2014) or with different antibiotic susceptibility (Novelli-Rousseau et al. 2018). It has also been used to discriminate between different *Acinetobacter* (Maquelin et al. 2006) or different *E. coli* strains (Jarvis and Goodacre 2004) amongst other examples. However, in these studies the expected phenotypes where known in advance. How to define what a phenotype is in a less-known system or a natural environment? In this work, we propose and validate the use of PhenoGraph –a tool derived from t-SNE and developed for the analysis of flow cytometry data-to do an automated identification of bacterial phenotypes.

The PhenoGraph algorithm was originally developed for mass cytometry data (Levine et al. 2015), a variation of flow cytometry which makes use of heavy metal ion tags instead of fluorochromes, resulting in more observed variables but at a lower acquistion speed (Spitzer and Nolan 2016). PhenoGraph was demonstrated to be highly effective for clustering purposes of single-cell Raman data, and returned a higher clustering performance compared to a more traditional hierarchical clustering approach. However, hierarchical clustering allows to inspect which cells are most similar to each other, a characteristic which is lost when using PhenoGraph. Therefore we want to reiterate that, as proposed by (Andrews and Hemberg 2018) for the analysis of single-cell RNA sequencing data, “[l]ikewise, no computational methods for dimensionality reduction, feature selection and unsupervised clustering will be optimal in all situations”. The algorithm of choice depends on the needs of the user. If a researcher wants to visualize subpopulations, we recommend the use of t-SNE. If identification of phenotypic populations is needed in an automated way, PhenoGraph is more appropriate. To assess which individual cells are phenotypically closest, hierarchical clustering can be used. Further investigation of the analysis of Raman data is needed, but investigating additional algorithms specifically developed for high-dimensional single-cell data might further support the impact of the use of Raman spectroscopy.

To improve the analysis speed of Raman spectroscopy, metallic substrates can enhance the signal (SERS), but also microfluidic chips (McIlvenna et al. 2016) or optical tweezers can be used (Xie, Chen, and Li 2005). An advantage of Raman spectroscopy is that it can be applied without the use of labels. This allows to analyse the biochemistry of samples even without knowing their nature. Raman spectroscopy also presents disadvantages. For instance, the Raman signals of certain compounds can be quite weak, making them difficult to detect or undetectable. The Raman signal of certain compounds can be composed of several peaks, or be unknown. Also, the background of samples can interfere with the Raman signal of bacteria. The equipment can be quite costly, depending on the type of Raman spectroscope.

Raman spectroscopy offers more parameters per cell compared to flow cytometry (hundreds versus typically three or four for microbial experiments). Thus, individual bacteria are described in a much larger multivariate space and can therefore be clustered into separate phenotypic populations. This explains why bacterial subpopulations can be visualized at the single-cell level using t-SNE (Fig. 3).The t-SNE results were confirmed with a PCA (Fig. S3).

The main downside of the use of label-free Raman spectroscopy is that the time of measurement is long: in this experiment, for single-cell label-free measurements, we used an acquisition time of 40 seconds per cell. Even when the acquisition time is lower –for instance, (Liu et al. 2016) reported a 1-3 seconds acquisition time to detect antibiotic susceptibility using surface-enhanced Raman spectroscopic biomarkers-the speed of Raman spectroscopy cannot match the high-throughput nature of flow cytometry for single-cell analysis.

### Raman spectroscopy allows to detect differences in biomolecules from one sample to another

Raman spectroscopy allows to detect biomolecules present in different phenotypic populations. Therefore, after automated identification of phenotypic populations, one can use the phenotypic groups to perform a variable selection strategy to select important regions in a data-driven way. We illustrated this approach using the Boruta algorithm, which was recently evaluated as one of the state-of-the-art variable selection methods using Random Forests for omics datasets (Degenhardt, Seifert, and Szymczak 2017). We found that a majority of selected spectral regions were the same according to treatment and automated phenotypic population identification using PhenoGraph (Fig. 3). This information can be used to infer how phenotypic populations are different at the level of their metabolism. To do so, we have based ourselves on a recent literature survey summarizing associations between Raman regions and certain biological compounds (Wang et al. 2016). The ten most important regions in function of phenotypic identification are listed in Table 2, along with the distribution of their intensities (Fig. 4). These regions correspond to carbohydrates and nucleic acids, as well as some unknown regions. An increase in the carbohydrate band (peaks 1042, 1046, 1050, 1057 cm-1) was observed for the stationary phase. The band at 1053 cm-1 could also be a nucleic acid peak, expected at 1054 cm-1. Nevertheless, these assignments for the Raman bands are tentative and based solely on a literature research, and thus proper validation of these results would have to be made in future experiments.

### How to define a phenotypic population?

In this work, we have steered microbial communities towards a certain growth stage, expecting that they would express a certain phenotype that could be retrieved using flow cytometry and Raman spectroscopy. However, in each one of these isogenic populations there might be subpopulations, as can be seen in Figure 2b. We acknowledge the difficulty in defining what a phenotypic population is, and setting a threshold to determine when one phenotypic population ends and another begins.

A similar problem exists in the area of bacterial taxonomy, where the similarity of 16S sequences is compared. In this case, an arbitrary threshold is set (e.g., 95% similarity at the genus level, 98.56% at the species level) (Stackebrandt and Goebel 1994; Kim et al. 2014). As explained by Beye et al. (2018), this cut-off was meant to standardize the use of 16S rRNA gene amplicon sequencing, but it had to evolve; i.e., the first threshold for the species level has changed from 97%, to 98.7%, to the current 98.65%. Even now, it is argued that these thresholds are not applicable to multiple genera (Mysara et al. 2017). In the case of phenotypic populations, we propose a definition based on their similarity (after setting a similarity threshold) and their ecology (their relationship with one another and with their environment). Quantifying their similarity can be done in a data-driven way, by means of for example clustering, at the resolution that is required for the specific research. This operational definition allows to define phenotypic populations depending on the research question, as long as researchers motivate and validate their choice. However, using this operational definition means that results cannot be compared across experiments or labs. This is why we highlight the need to find a more standard way to define ‘basic phenotypic units’, that would allow to measure phenotypic traits and determine if bacteria belong to the same phenotypic population.

We propose to use algorithms -such as hierarchical clustering, t-SNE or PhenoGraph, applied throughout this paper-to define, visualize and characterize phenotypic populations. t-SNE is a well-known technique to visualize high-dimensional single-cell data, being commonly applied to visualize for example cytometry and single-cell RNA sequencing data (Amir et al. 2013; Andrews and Hemberg 2018). Our results confirm that it can be used as an ‘off-the-shelf’ visualization method to detect phenotypic populations in Raman data when applied to microorganisms.

Bacteria were grown in 9 different conditions (three replicate cultures of three growth stage conditions) to steer the same *E. coli* population to a different morphological and/or metabolic state –to steer them into 9 phenotypic populations. While hierarchichal clustering was able to find eight of these phenotypic populations, PhenoGraph was able to retrieve all nine of them, resulting in a higher ARI as well.

t-SNE and PhenoGraph were also applied to an external dataset from Teng et al., 2016, consisting of *E. coli* that had been treated with different agents, and measured at several time points. We showed that PhenoGraph was capable of differentiating the time points per treatment. Interestingly, two subpopulations were identified per treatment, although samples were measured in triplicate. These corresponded to two replicates, where the third was either assigned to one subpopulation or divided between both (Fig. 6, Fig. S6). Our group has previously shown how small technical variations can create subpopulations that have no biological meaning (García-Timermans et al. 2018), which might explain these findings.

Each algorithm has its own advantages and disadvantages. t-SNE is a highly effective technique to visualize high-dimensional single-cell data, which we confirmed for the Raman analysis of microorganisms. However, automated clustering of the data is not possible using t-SNE without additional algorithms, such as PhenoGraph.

### Conclusions

The results of this research suggest that:

- Flow cytometry is a more high-throughput technology than label-free Raman spectroscopy, but Raman describes bacterial cells in many more variables, without the need for staining.
- Flow cytometry can be applied to quantify differences in phenotypic heterogeneity at the population level, whereas Raman spectroscopy has sufficient resolving power to identify separated phenotypic populations at the single-cell level.
- Raman spectroscopy provides the possibility to infer which metabolic properties define different phenotypic populations, and potentially exploit this information for bioprocess monitoring.
- We propose a workflow to automatically identify bacterial phenotypes, based on Raman spectral data. We also recommend t-SNE to visualize Raman data.
From a broader perspective, one can motivate that phenotypic populations depend on a similarity threshold, which can be set in clustering algorithms, and on their ecological niche. We therefore suggest that researchers try to include validation controls in their experimental setup, in order to motivate the threshold of choice in the algorithm.

## Materials & Methods

### Cell culture

To determine the growth stages of the cell culture (lag, log and stationary phase), *Escherichia coli* DSM 2092 was grown in nutrient broth (NB, Oxoid, United Kingdom) at 28°C, 120 rpm shaking in three replicates. Cultures had an initial concentration of 10^6^ cells/ml, measured with a BD Accuri C6 flow cytometer (BD Biosciences), following the protocol from Van Nevel 2013. The samples were incubated in the dark for 30 h at 28°C, during which optical density (OD, λ = 620 nm) measurements were automatically collected each hour using a microtiter plate reader (Tecan Infinite M200 Pro; Tecan UK, Reading, United Kingdom). The growth phases were assigned after fitting the results with the function *SummarizeGrowth()* from the ‘Growthcurver v0.30’ R package (Sprouffske and Wagner 2016). Cells were harvested 1h, 7h30 and 24h after inoculation, corresponding accordingly to the lag, log and stationary phases of *E.coli* (see Fig. S1). Nutrient broth was included as a negative control.

### Sample preparation

Samples were measured immediately in the flow cytometer after sampling. For Raman spectroscopy, samples were harvested and fixed in formaldehyde 4% (Sigma-Aldrich) dissolved in PBS (protocol from Bio-Techno Ltd., Belgium) following the protocol from (García-Timermans et al. 2018). First, 1mL of the cell suspension was centrifuged for 5 min at room temperature and 1,957 × *g*. For the samples in the lag phase, up to 10 mL were suspended until a pellet could be seen. The supernatant was discarded and cells were suspended in filtered and cold PBS (4°C). The samples were again centrifuged at 1,957 × *g* for 5 min at room temperature. The supernatant was discarded and the pellet was re-suspended in filtered formaldehyde 4%. The cells were allowed to fix for 1h at room temperature (21°C). Then, the samples were centrifuged at 1,957 × *g* for 5 min at room temperature and washed twice with cold PBS (4°C). Cells were stored at 4°C and analysed in the Raman spectroscope within the week.

### Flow cytometry

Fresh samples taken at the lag, log and stationary phase were diluted in filtered PBS and stained with SYBR Green I 1% (Thermo Fisher) during 13 min at 37°C. They were measured with the flow cytometer BD Accuri C6 (BD Biosciences). This resulted in a multivariate description of each cell by four fluorescence detectors (FL1: 533/30 nm, FL2: 585/40 nm, FL3: > 670 nm long pass, FL4: 675/25 nm), of which the FL1 detector was targeted by SYBR Green I, and two scatter detectors (forward scatter, FSC and side scatter, SSC). The channels FSC-H, SSC-H, FL1-H and FL3-H were used for data analysis.

#### Single-cell analysis

##### t-distributed stochastic neighborhood embedding (t-SNE)

t-SNE is a dimensionality reduction technique developed for the visualization of high-dimensional data (Van Der Maaten and Hinton 2008). The *TSNE()* function from the scikit-learn machine learning library was used (Pedregosa et al. 2011, v0.19.1). Principal component analysis was set as initialization method. TSNE was run with default settings unless reported otherwise. Data were first transformed by the function f(x) = asinh(x), and next normalized so that each channel has a mean of zero and standard deviation of one.

##### Principal component analysis (PCA)

Flow cytometric single-cell data were analysed with the *PCA()* function from the scikit-learn machine learning library after normalization. Data were first transformed by the function f(x) = asinh(x), and next normalized so that each channel has a mean of zero and standard deviation of one.

##### Community analysis

The PhenoFlow R package (Props et al., 2016) was used for the analysis. Four channels (FL1-H, FL3-H, FSC-H and SSC-H) were selected to derive a phenotypic fingerprint for each sample. Bacteria were gated to differentiate from background noise as shown in Fig. S2. As quality control, the stability of the FL1 signal over time was checked. A 128 x 128 binning grid was constructed for each pairwise combination of these channels (resulting in 6 in total). Next, a bin a kernel density estimation was performed to determine the density per bin (with a Gaussian kernel density bandwidth of 0.01). Then, all bins are concatenated to a one-dimensional vector, representing the cytometric fingerprint. Data were normalizedtransformed usingby the function f(x) = asinh(x) transformation. At least 10.000 cells were measured per sample.

##### Principal component analysis (PCA) and principal component analysis (PCoA)

The pulled information for every group was analysed with the function *fviz_pca_ind()* from the R package ‘factoextra’ (Kassambara and Mundt 2017).

Principal coordinate analysis (PCoA, also known as multidimensional scaling) was calculated based on the Bray-Curtis dissimilarities between all fingerprints. The function *beta_div_fcm()* from the R package ‘PhenoFlow’ was used (Props et al. 2016).

### Raman spectroscopy

Fixed samples were centrifuged at 1,957 x g for 5 min at room temperature and re-suspended in cold Milli-Q water (Merck-Millipore) (4°C). Then, a 5μL drop was allowed to dry until evaporation on a CaF2 slide (grade 13 mm diameter by 0.5 mm polished disc, Crystran Ltd). As control for the instrument performance, a silica gel was measured with a grating of 600 – mm/g, with a 1 second time exposure and 10 accumulations. Laser power was also monitored to detect possible variations. Bacteria were measured with a grating of 300 –mm/g, with a 40 second exposure time and 1 accumulation. More information on the Raman spectroscope and data collection is included in the Raman aid (see Table 3). The metadata were collected following the guidelines from (García-Timermans et al. 2018) and can be found in the Supplementary Information.

#### Raman spectra pre-processing

The Raman spectra were analysed in the 600-1800 cm^-1^ region, and baseline correction using the SNIP algorithm (ten iterations) and normalization were performed. The area under the curve (AUC) normalization was calculated with the MALDIquant package (v1.16.2) (Gibb and Strimmer 2012).

#### Single-cell analysis

##### t-distributed stochastic neighborhood embedding (t-SNE)

Raman single-cell data was analyze dusing t-SNE. The *TSNE()* function from the scikit-learn machine learning library was used. Principal component analysis was set as initialization method. TSNE was run with default settings unless reported otherwise. Each region in the spectra was normalized to have zero mean and standard deviation of one.

##### Principal component analysis (PCA)

Single-cell Raman spectra were analysed with the function fviz_pca_ind() from the R package ‘factoextra’ (Kassambara and Mundt 2017) or with the PCA() function from the scikit-learn machine learning library after normalization of the spectra, so that each region has a mean of zero and standard deviation of one.

##### Hierarchical clustering

To measure how dissimilar the samples were, we calculated the spectral contrast angle (Wan, Vidavsky, and Gross 2002) between individual cells based on Raman spectra. Then, clusters were determined in an agglomerative way, through Ward’s method (ward.D2) from the *fastcluster* R package (Müllner 2018). Hierarchical clustering was implemented using the *hclust()* function from the stats package (R Core Team 2018).

##### PhenoGraph

PhenoGraph is a clustering algorithm specifically designed for the analysis of high-dimensional flow- or mass-cytometry data (Levine et al. 2015). It employs a two-step approach, in which for every cell its *k*-nearest cells of similar phenotypic populations are grouped together. This means that, if *N* denotes the number of cells, *N* neighbourhoods are created. Next, a weighted graph is created on these sets of cells. The weight between nodes scales with the number of neighbours that are shared. The Louvain community detection method is implemented to cluster the graph by maximizing the modularity of different groupings of the nodes (Blondel et al. 2008). The PhenoGraph algorithm was run with default settings, in which k was evaluated for different values between five and 100 (github.com/jacoblevine/PhenoGraph). PhenoGraph was run after normalization of the spectra, to have zero mean and standard deviation of one.

##### Adjusted Rand Index

Clustering results from both hierarchical clustering and PhenoGraph were quantified by the Adjusted Rand Index (ARI) (Hubert and Arabie 1985). The ARI was calculated with the *adjusted_rand_score()* function from the scikit-learn machine learning library (v0.19.1) (Pedregosa et al. 2011). The Rand index is defined as the number of pairs of instances that are in the same group or in different groups based on two partitions, which is divided by the total number of pairs of instances. This index is then corrected for the expected index, which is based on random clustering in which the elements per cluster are shuffled between clusters. A value of 1 resembles the perfect match between cluster assignments and ground truth labels, a value of 0 resembles random clustering and a negative value (up to -1) resembles arbitrarily worse clustering.

##### Boruta variable selection

The Boruta variable selection extends on traditional variable selection using Random Forest based variable importance measures. The method includes shadow variables, which are copies of original variables that have been permutated. By performing a Random Forest analysis multiple times, one can decide by means of multiple hypothesis testing which variables are relevant with a certain significance level compared to the most relevant shadow variable (Kursa and Rudnicki 2010). It has been recently proposed as one of the most accurate and stable variable selection methods based on Random Forest based (Degenhardt, Seifert, and Szymczak 2017). The Boruta algorithm from the *Boruta* R package was run, using the default settings (v6.0.0) (Kursa and Rudnicki 2010).

##### Statistical test on Boruta outcome

The ten most relevant regions for classification according to the Boruta algorithm were selected. The intensity of these peaks amongst the growth phases were compared with the Wilcoxon rank sum test with a Benjamini-Hochberg correction (upon rejection of the null hypothesis). The functions pairwise.wilcox.test() and p.adjust() from the R package stats v3.5.1 (R Core Team 2018) were used.

#### Data availability

Data and code to reproduce analysis is available on the following repository: https://github.com/CMET-Ugent/FCMvsRaman.

Data analysis was conducted using the program R (R Core Team 2018), RStudio (RStudio team and RStudio 2016) and Python.

#### External dataset

We included the dataset from (Teng et al. 2016) in order to validate the generalizability of the PhenoGraph and t-SNE algorithms for the analysis of label-free bacterial Raman data. As described in their article, they tested the stress response of *E. coli* to six chemical stressors at different time intervals with label-free Raman spectroscopy: ethanol, antibiotics ampicillin and kanamycin, n-butanol or heavy metals Cu^2^+ (CuSO4) and Cr^6^+ (K2CrO4). Teng et al. showed that each of these treatments resulted in a different phenotype. In other words, each treatment resulted in a unique Raman characterization of cells, which should group together upon analysis. These treatments were therefore used as label according to which PhenoGraph or t-SNE should group the cells. Three biological replicates of the cell culture were made, and 20 cells were tested per replicate. Bacteria were sampled at different stages of the cell growth. The Raman spectra of the stressed cells were collected after the treatment (5, 10, 20, 30 and 60 min, 3h and 5h).

##### Data availability

Our code and data to reproduce the analysis is available at https://github.com/CMET-Ugent/FCMvsRaman.

## Author contributions

CGT and PR co-wrote the paper with contributions from JH, FMK, RP, AS, WW, and NB. CGT collected the data. PR, CGT and RP performed the data analysis. CGT, PR, JH, FMK, RP, WW and NB designed the study. All authors read and approved the final version of the manuscript.

## Acknowledgements

The authors would like to thank the funding that made possible this research. CG is funded by Qindao Beibao Marine Science & Technology Co. Ltd., Qingdao West-coast economic new area, China. PR is funded by Special Research Fund (BOFSTA2015000501) from Ghent University. RP is funded by by Ghent University (BOFDOC2015000601). JH is funded by the Flemish Fund for Scientific research (FWO-Vlaanderen, 1S80618N). AGS acknowledges support of BOF UGent (BOF14/IOP/003, BAS094-18, 01IO3618) and FWO-Vlaanderen (G043219). This work was supported through the Geconcerteerde Onderzoeksactie (GOA) from Ghent University (BOF15/GOA/006) and the MICCAS project (project grant no. 3G020119 of the FWO Flanders). The authors would also like to thank Dmitry Khalenkow for his help setting up the Raman microscope.

---

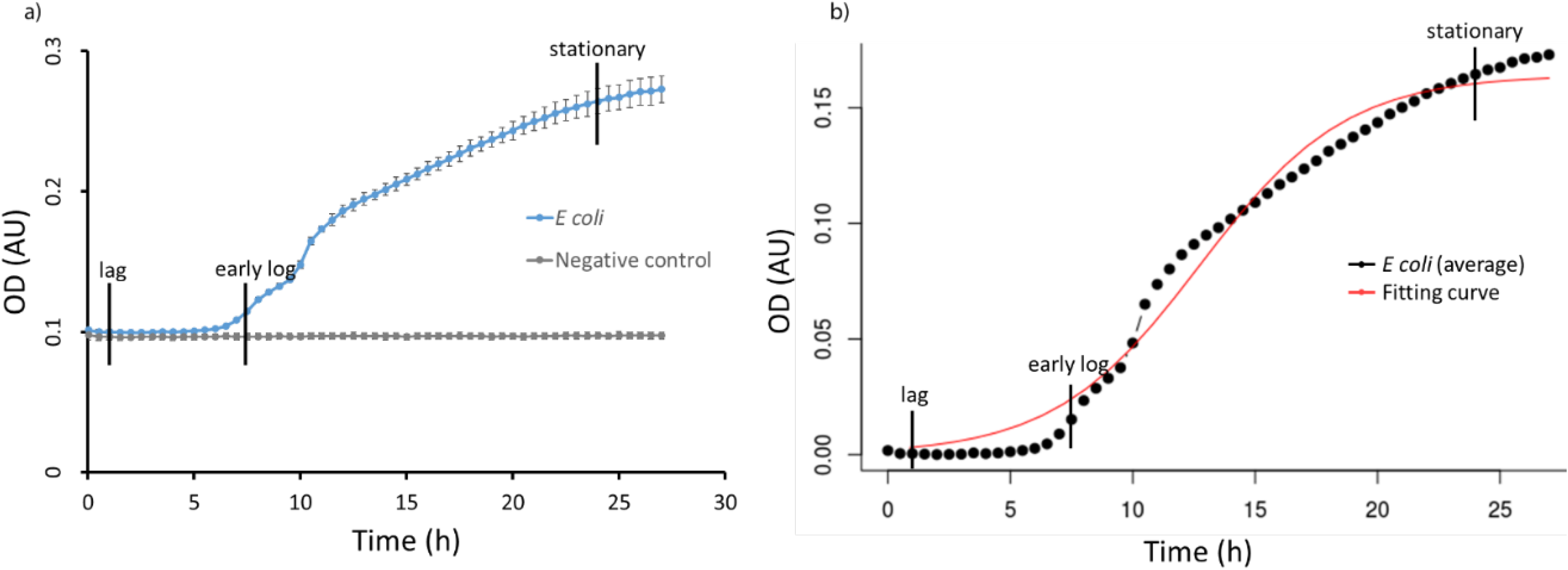

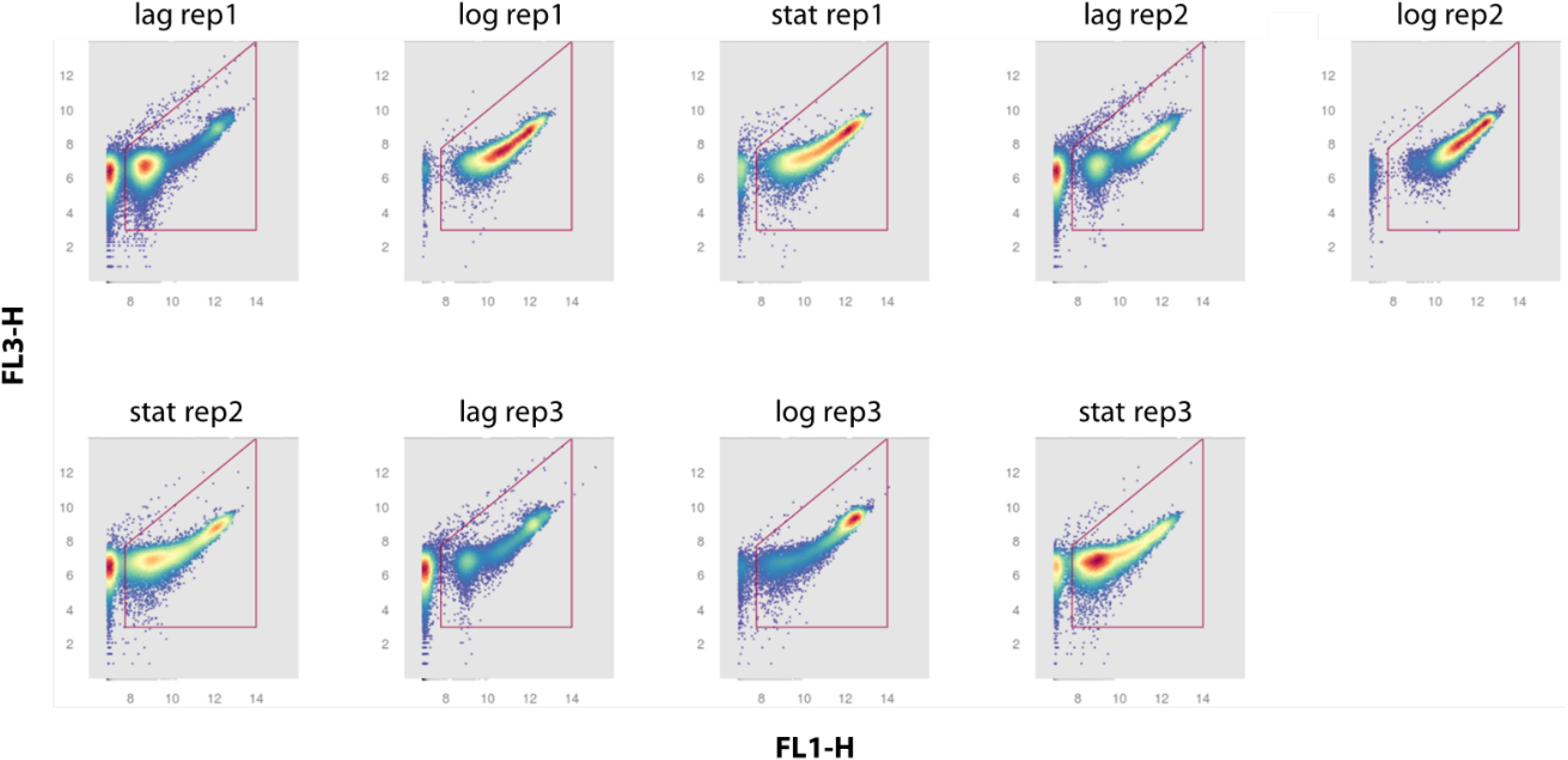

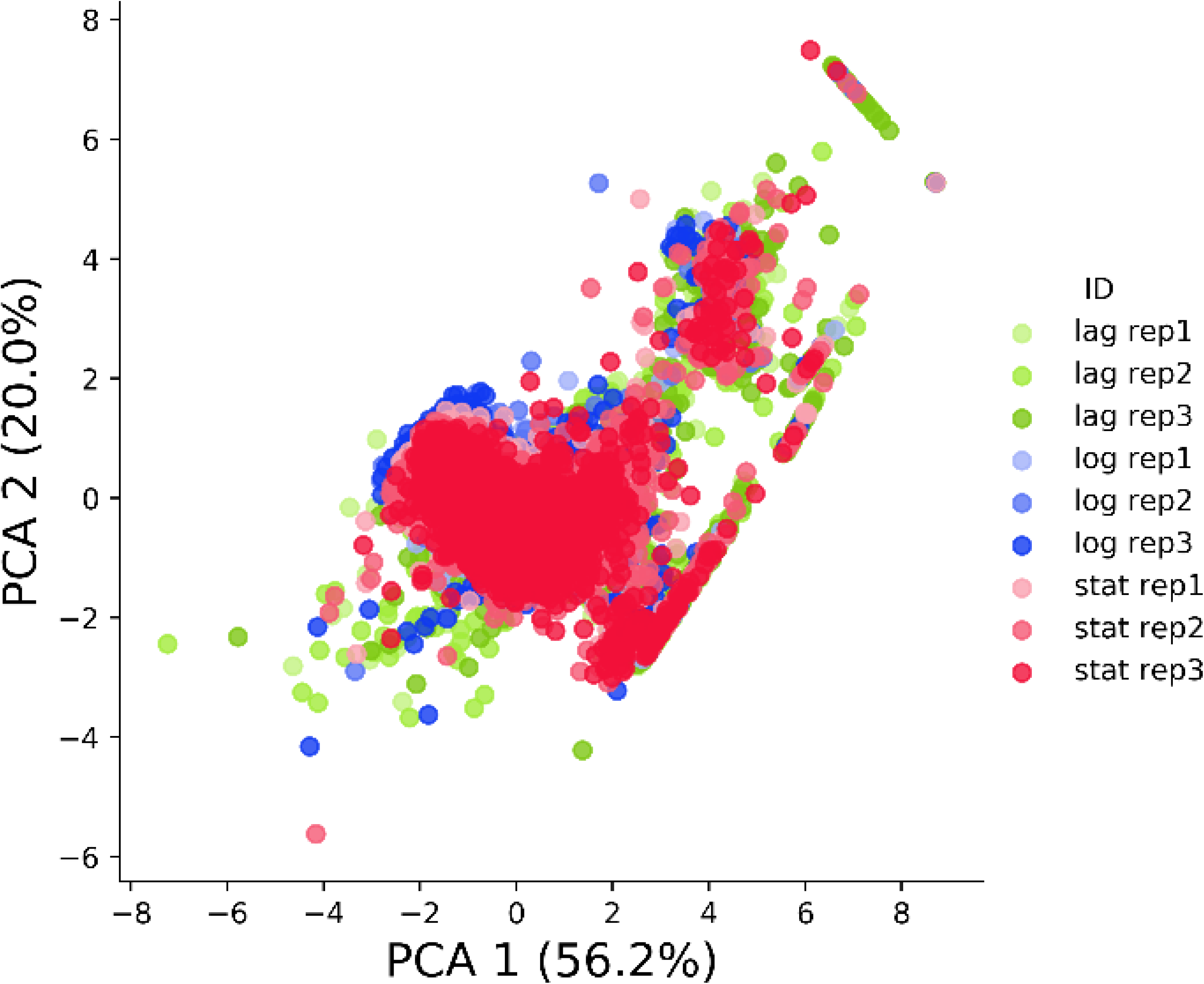

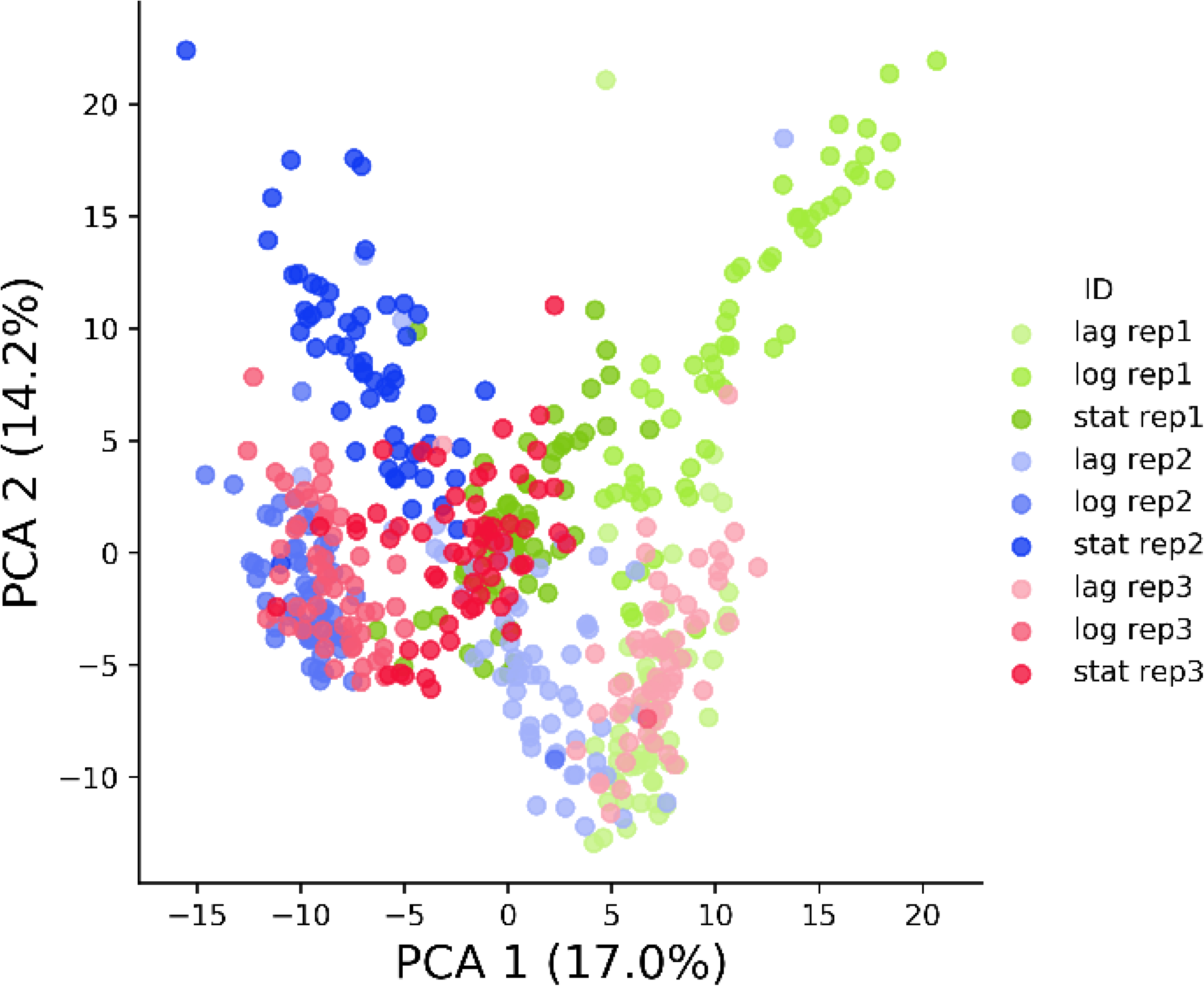

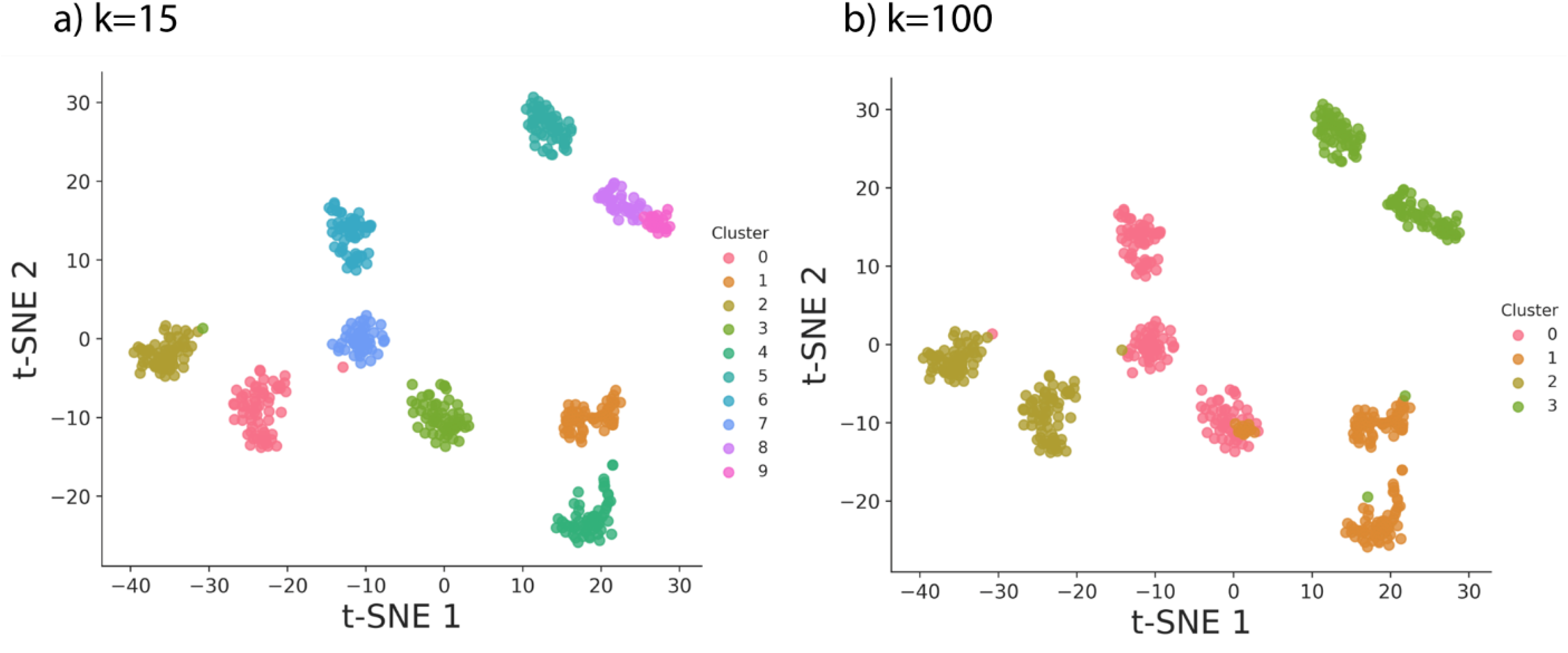

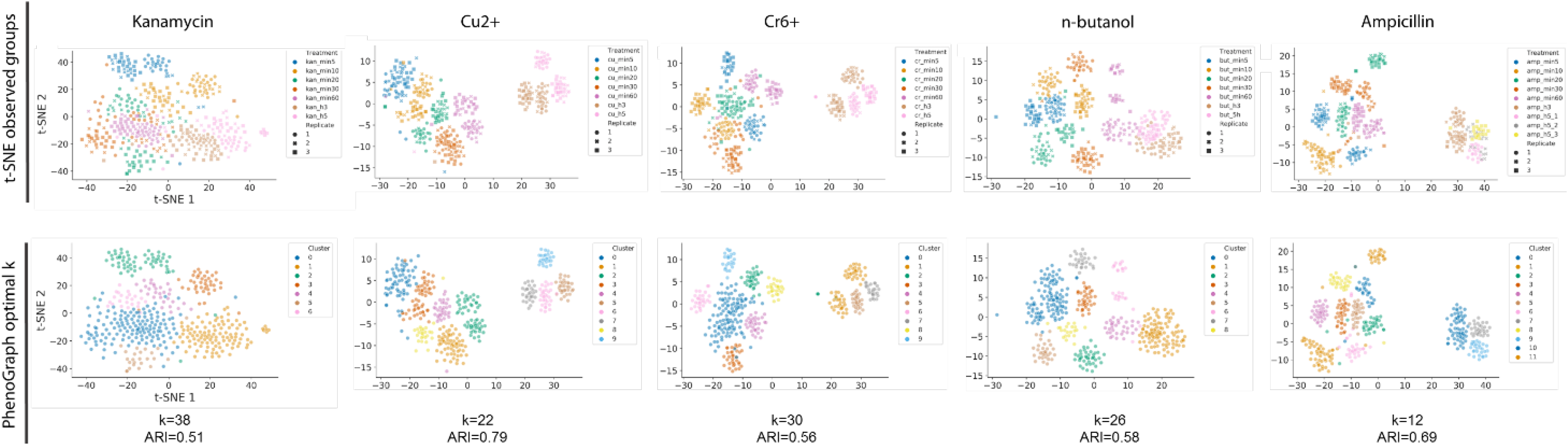

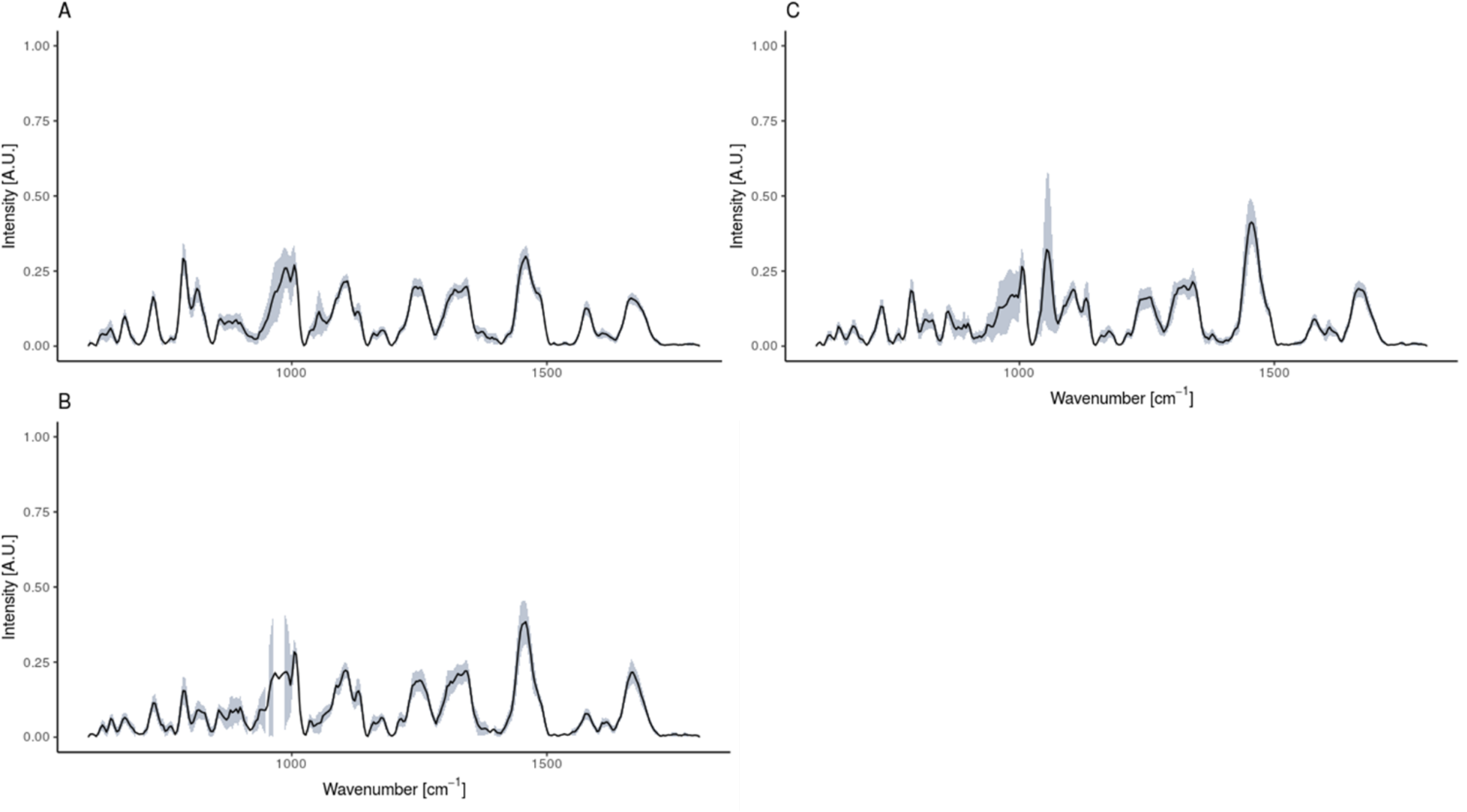

