## Supplementary Information for "Discriminating bacterial phenotypes at the population and single-cell level: a comparison of flow cytometry and Raman spectroscopy fingerprinting"

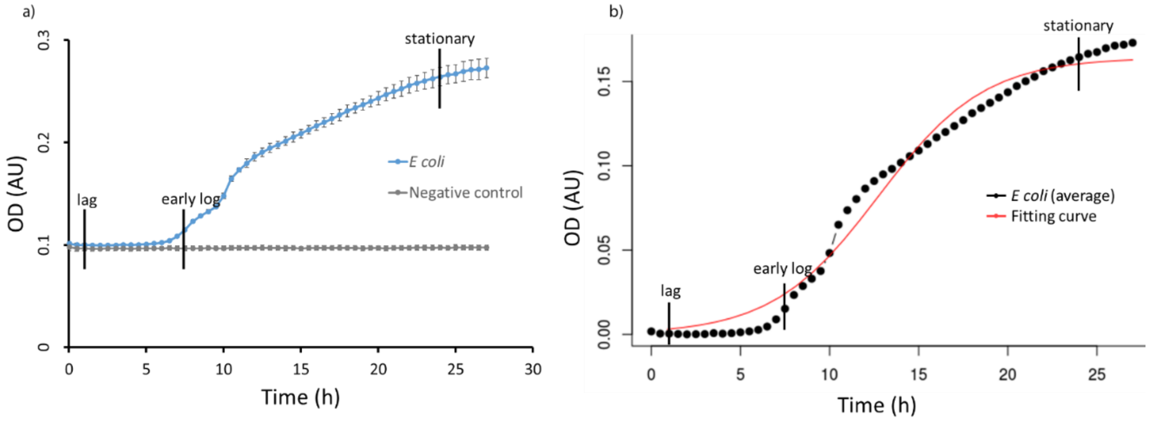


Fig S1: Growth curve for *E. coli* in nutrient broth at 28°C, 120 rpm shaking. Three replicates of the cell culture were made. (a) OD results. In blue, the results for the E. coli; in grey, the negative control (medium). (b) Fitting model for the *E. coli* OD results (after background substraction) to assign the lag, log and stationary phases.

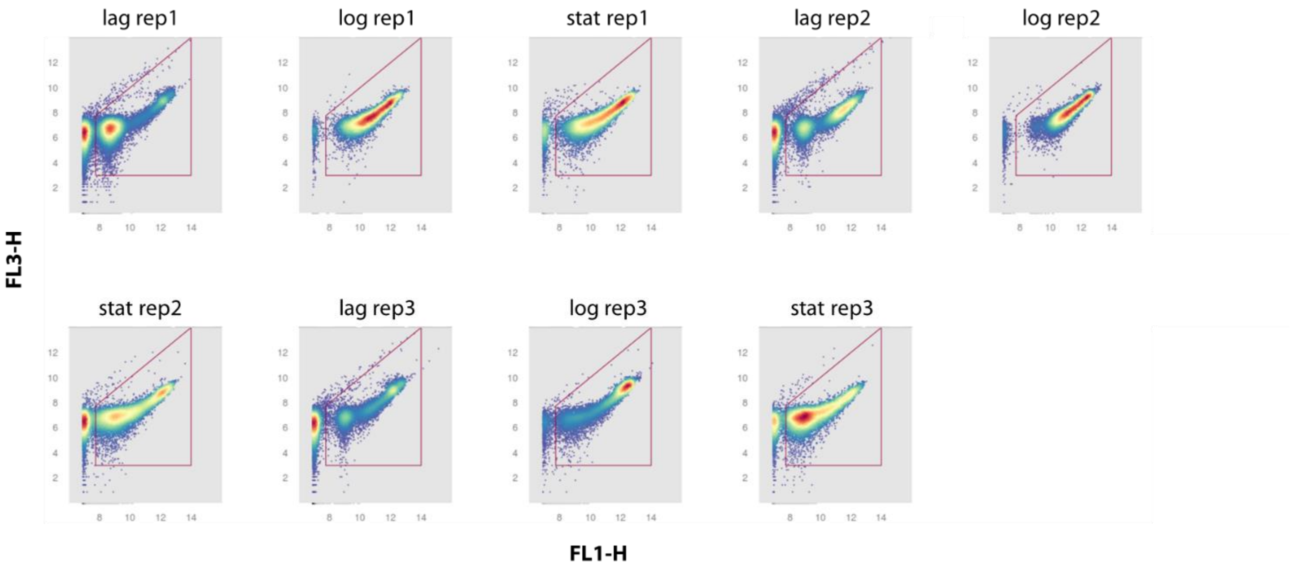


Fig S2: Gating strategy for the flow cytometric data. Arcsinh transformed data of FL1/FL3 in a density plot (see Materials & Methods).

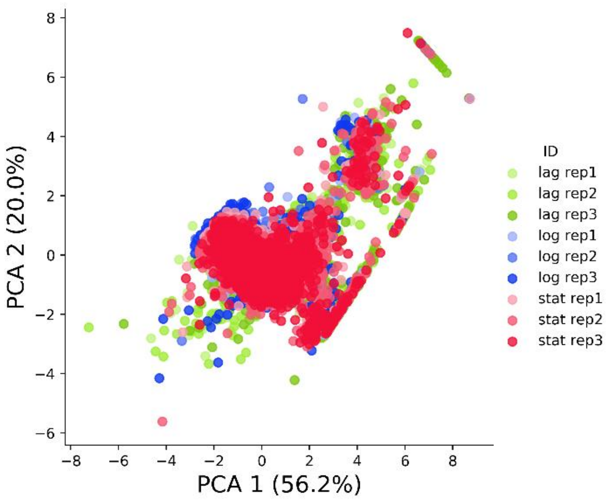


Fig S3: First two components of a PCA for single-cell flow cytometric data. Colors correspond to growth phases, shapes to replicates.

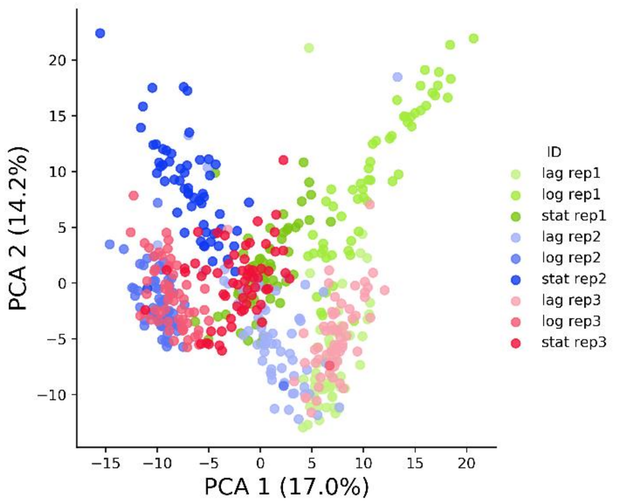


Fig S4: First two components of the PCA for Raman spectra of individual cells. Colors correspond to growth phases, shapes to replicates.

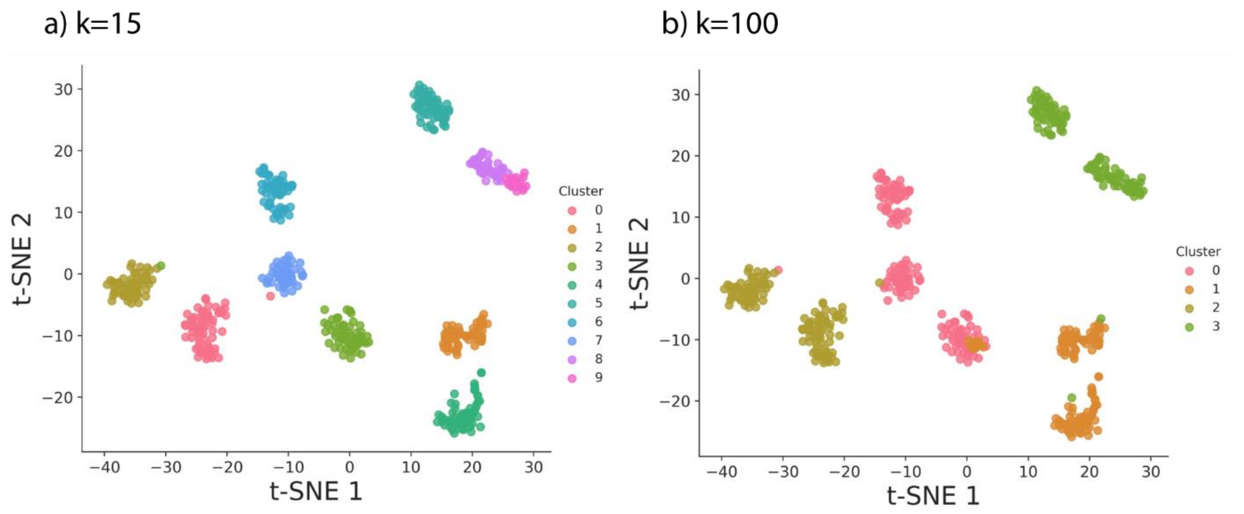


Fig S5: Visualization of PhenoGraph clustering results of the Raman data derived from the **E. coli** culture presented in Figure 3 for different k: (a) k=15, (b): k=100.

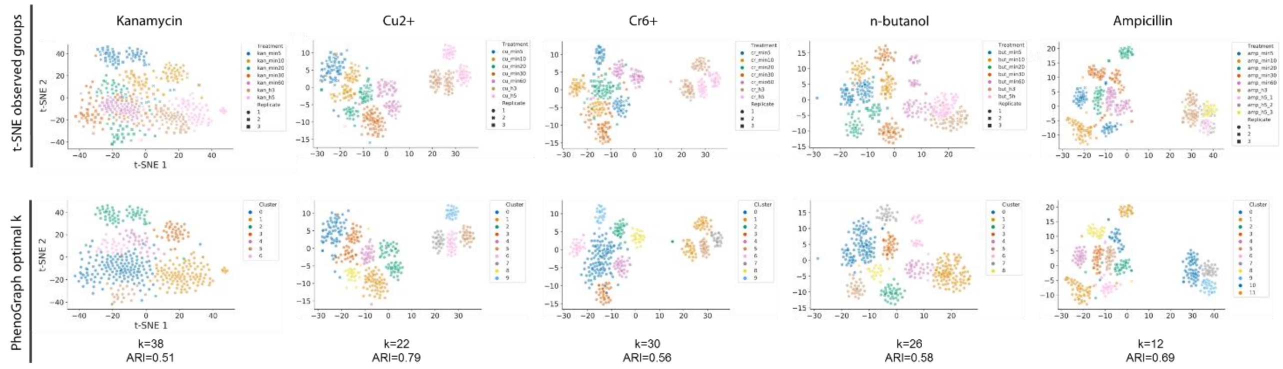


Fig. S6: Ramanone results for all the groups. On top, the t-SNE for the observed phenotypic groups. At the bottom, the PhenoGraph results after calculating the optimal number of clusters (highest ARI).

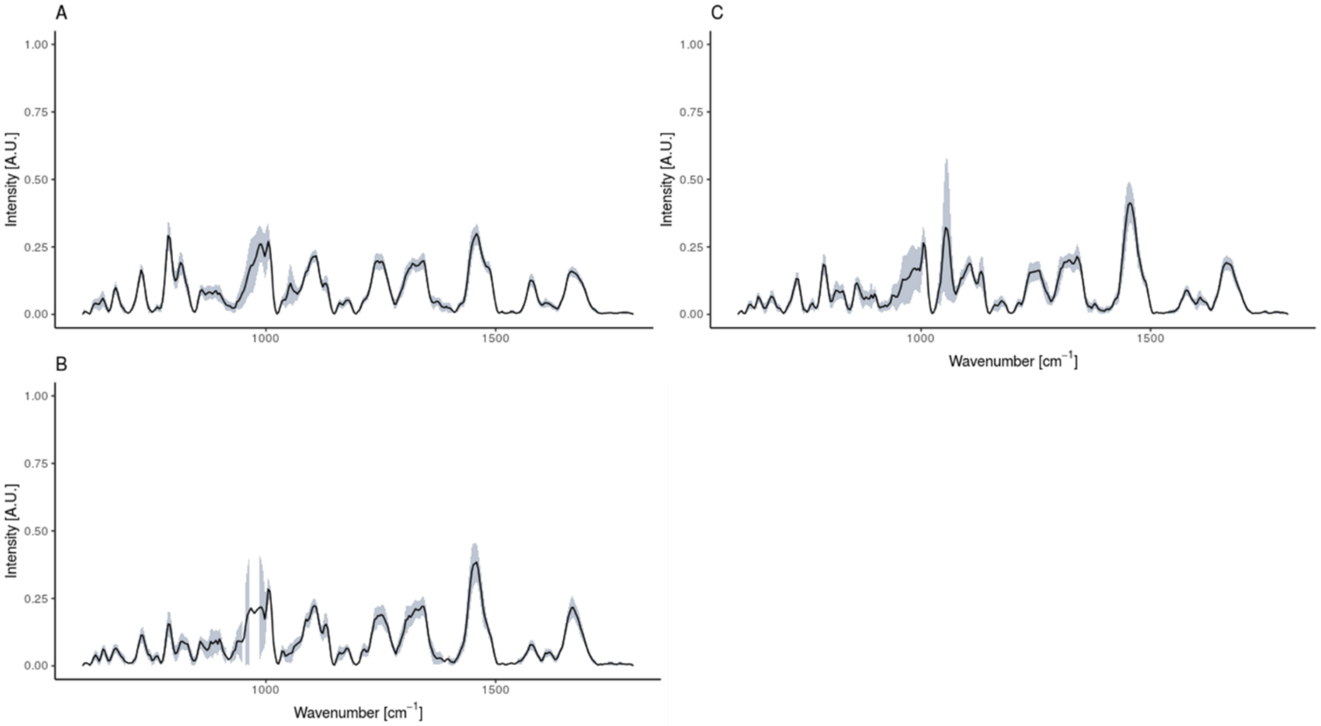


Fig. S7: Raman spectra of the cells according to growth stage. *E. coli* harvested in the A) lag B) log and C) stationary phase. In lighter color, the standard deviation is represented

| **Experiment overview:** | |
|---|---|
| Hypothesis | *E. coli* in different growth stages and/or cells from different replicates will differ in their Raman spectra |
| Variable(s) tested | *E.coli* were fixed in log, lag and stationary phase. Three replicates of the cell culture. |
| Conclusions | Differences in the Raman spectra were found when cells differed in the growth stage and/or belonged to another culture |
| Quality control (internal/external) | Silica gel check |
| **Samples and sample acquisition** | |
| Material and source | *Escherichia coli DSM 2092* |
| Growing conditions/sampling | Cells were grown in LB, 25°C, 120 rpm shaking |
| Filename format: <Replicate number>_<Treatment>_<Cell number> | |
| Label in the samples | No label used |
| Fixation method | Filtered PFA 4% |
| Integration time | 40 sec |

| | |
|---|---|
| Accumulations | 1 |
| Grid | 300 –mm/g |
| **Instrument** | |
| Laser | 785 nm excitation diode laser (Toptica). 175 mW of power before the objective. |
| Quality control | A silica gel sample was measured with a grating of 600 –mm/g, with a 1 second time exposure and 10 accumulations. Laser power was also monitored to detect possible variations. |
| Objective used (magnification ) / Numeric aperture (NA) | 100x/0.9 NA (Nikon) |
| Camera | -70 °C cooled CCD camera (iDus 401 BR-DD, ANDOR) |
| Dry/water/oil objective | Dried samples |
| Model of spectroscope | WITec Alpha300R+ |
| Other specifications (chromatic/flat field correction/other) | |
| **Data analysis** | |
| Background subtraction method (if used) | No. Measurements with cosmic rays were deleted |
| Normalization method (peak /min-max /area under-curve /other) | Area under the curve (‘Total Ion Count’) |
| Smoothing and interpolation (if done) | Baseline correction |
| Statistics/Machine learning algorithm | ‘MicroRaman’ package (GitHub). Spectral contrast angle, ward.D2 dissimilarity and hierarchical clustering. Boruta |
| Accessibility | https://github.com/CMET-Ugent/FCMvsRaman. |
| Other relevant information | |

Table S1: Metadata aid for Raman spectra
